# In Situ Replication Rates for Uncultivated Bacteria in Microbial Communities

**DOI:** 10.1101/057992

**Authors:** Christopher T. Brown, Matthew R. Olm, Brian C. Thomas, Jillian F. Banfield

**Author notes:** Address: McCone Hall, Berkeley, CA 94720. **Author Contributions:** CTB and JFB developed the iRep and bPTR methods. MRO ordered and oriented draft genome sequences for bPTR calculations and conducted kPTR analyses. CTB conducted the iRep, bPTR, and kPTR comparisons, and determined the accuracy of the iRep method. JFB binned the adult human metagenome and curated the Deltaproteobacterium genome, with input from CTB. CTB implemented the iRep method. BCT provided bioinformatics support. CTB and JFB drafted the manuscript. All authors contributed to iRep development, reviewed results, and approved the manuscript.

## Abstract

Culture-independent microbiome studies have revolutionized our understanding of the complexity and metabolic potential of microbial communities, but information about in situ growth rates has been lacking. Here, we show that bacterial replication rates can be determined using genome-resolved metagenomics without requirement for complete genome sequences. In human infants, we detected elevated microbial replication rates following administration of antibiotics, and bacterial growth rate anomalies prior to the onset of necrotizing enterocolitis. We studied microorganisms in subsurface communities and determined that a diverse group of groundwater-associated bacteria typically exhibit slow growth rates, despite significant changes in geochemical conditions. All microbiome studies will be advanced by measurements of replication rates that can identify actively growing populations, track organism responses to changing conditions, and provide growth rate information needed for modeling.

## Main Text

Dividing cells in a natural population contain, on average, more than one copy of their genome (Fig. 1). In an unsynchronized population of growing bacteria, cells have replicated their genomes to different extents, resulting in a gradual decrease in average genome copy number from the origin to terminus of replication^1^. This can be detected by measuring changes in DNA sequencing coverage across complete genomes^2^. Because bacteria replicate their genomes bidirectionally from a single origin of replication^3,4^, the origin and terminus of replication can be determined based on this coverage pattern^2^. Further, early studies of bacteria cultures determined that cells can achieve faster divisions by simultaneously initiating multiple rounds of genome replication^5^. This leads to an average of more than two copies of the genome in very rapidly growing cells.

**Figure 1.**
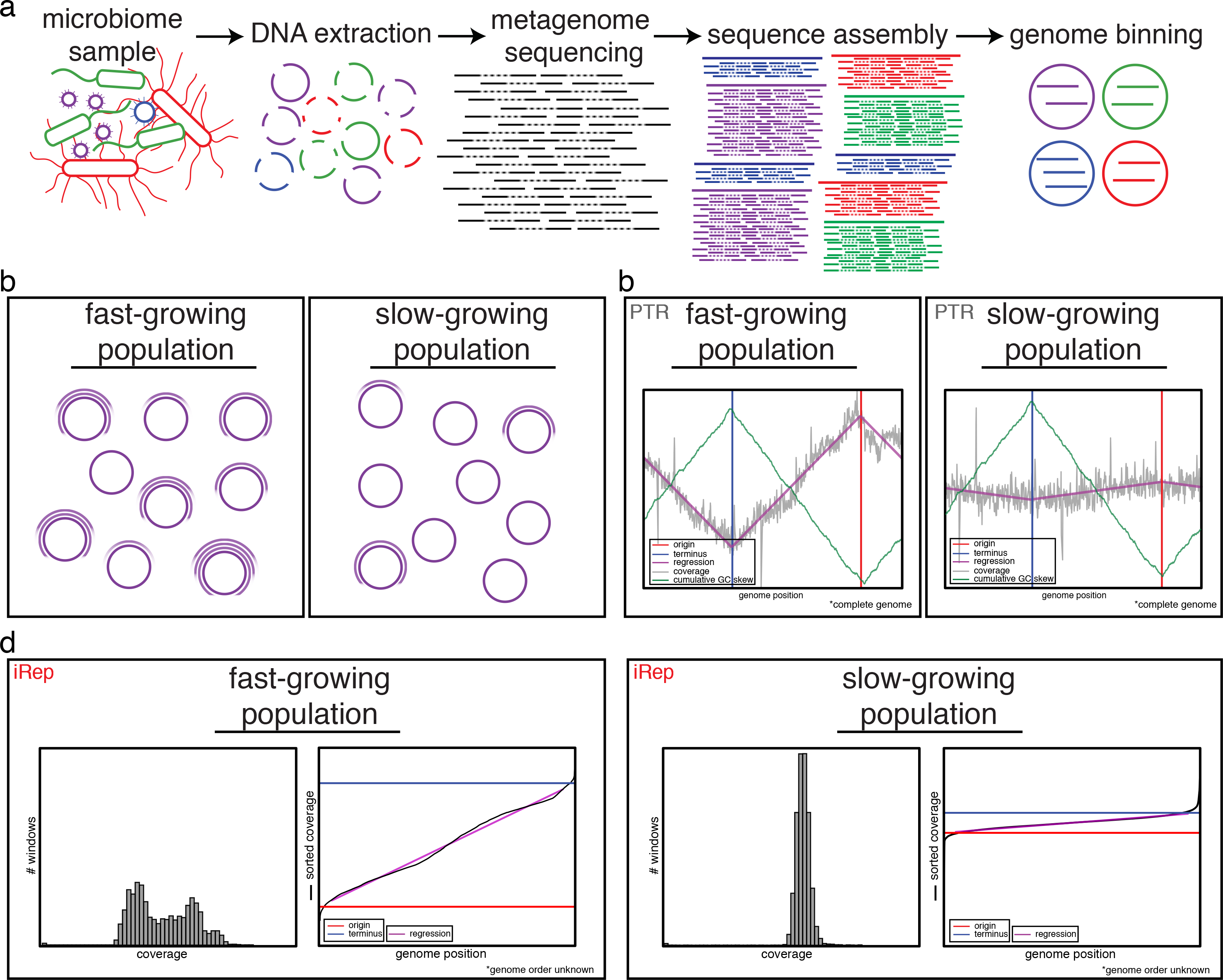
iRep determines growth rates for bacteria using genome-resolved metagenomics. (**a**) Genome-resolved metagenomics involves DNA extraction from a microbiome sample followed by DNA sequencing, assembly, and genome binning. Binning is the grouping together of assembled genome fragments that originated from the same genome. This can be done based on shared characteristics of each fragment, such as sequence composition, taxonomic affiliation, or abundance. (**b**) Populations of bacteria undergoing rapid cell division differ from slowly growing populations in that the individual cells of a growing population are more actively in the process of replicating their genomes (purple circles). (**c**) Differences in genome copy number across a population of replicating cells can be determined based on sequencing read coverage over complete genome sequences. The ratio between the coverage at the origin (“peak”) and terminus (“trough”) of replication (PTR) relates to the growth rate of the population. The origin and terminus can be determined based on cumulative GC skew. (**d**) If no complete genome sequence is available, it is possible to calculate the growth rate based on the distribution of coverage values across a draft-quality genome using the iRep method. Coverage is first calculated across overlapping segments of genome fragments. Growing populations will have a wider distribution of coverage values compared with stable populations (histograms). These values are ordered from lowest to highest, and linear regression is used to evaluate the coverage distribution across the genome in order to determine the coverage values associated with the origin and terminus of replication. iRep is calculated as the ratio of these values.

Korem *et al.* showed that the ratio of sequencing coverage at the origin compared to the terminus of replication could be used to measure replication rates for bacteria in cultures and in the mouse gut^6^. Because the origin and terminus correspond with coverage peaks and troughs, respectively, the authors refer to the method as PTR (peak-to-trough ratio). They also applied this method to calculate replication rates for some organisms associated with the human microbiome. However, the requirement of this method for mapping of sequencing reads to complete (closed and circularized) reference genomes is a major limitation.

Metagenomics methods routinely generate draft genomes for organisms for which no complete genomes are available^7–14^ (Fig. 1). Often these organisms are from previously little known and unknown microbial phyla, and thus the genomes are vastly different from the complete genomes in public databases^12–18^. In some cases, hundreds or even thousands of draft or near-complete genomes can be recovered from a single ecosystem. Thus, there is clear motivation to extend coverage-based replication rate analysis to enable measurements based on these genomes. Here, we introduce a novel method that can be applied to draft genomes to measure replication rates based on sequencing coverage trends. The method works, despite the fact the order of the fragments is unknown. This will enable the analysis of replication rates for large numbers of bacteria in microbial community context. Unlike PTR, this approach will find applications in virtually every natural and engineered ecosystem, including complex systems such as soil from which representative complete genomes for the vast majority of bacteria are not available.

## Results

### Calculating the Index of Replication (iRep) from genome-resolved metagenomics

The method that we developed is based on the well-established phenomenon that DNA sequencing coverage for replicating bacterial populations decreases smoothly from the origin to terminus of replication, and that this difference in coverage is proportional to growth rates. However, rather than identify these regions and coverage values directly, our method evaluates the total change in coverage across genome fragments. As we show below, these values are comparable to PTR, but since they are derived differently we refer to this metric as the Index of Replication (iRep).

iRep values are calculated by mapping metagenome sequencing reads to the collection of assembled sequences that represent a draft genome (Fig. 1; **see Online Methods and Code Availability**). The read coverage is evaluated across each scaffold over sliding windows. Data from different fragments are combined, and then the values are ordered from lowest to highest coverage to assess the coverage trend across the genome. Because coverage windows are rearranged in the process, the order of the fragments in the complete genome need not be known. Extreme high and low coverage windows are excluded, as they often correlate with highly conserved regions, strain variation, or integrated phage. Finally, the overall slope of coverage across the genome is used to calculate iRep, a measure of the average genome copy number across a population of cells.

### iRep measurements are accurate for complete and draft-quality genomes

In order to evaluate the ability of the iRep method to measure replication rates, we compared iRep to PTR using data from the growth rate experiments used by Korem *et al.* to establish the PTR method^6^. Since there is no open-source version of their software, we re-implemented the PTR method, with some significant modifications that include an option to determine the origin and terminus positions based on GC skew^19^ (**see Online Methods and Code Availability**). PTRs generated using the Korem *et al.* software (kPTRs) use a genome database of unknown composition that cannot be modified, and no metrics for evaluating measurement reliability are provided. These limitations are addressed in our new PTR implementation (bPTR). Calculated kPTRs and bPTRs were highly correlated, and each was correlated with iRep (Fig. 2a and **Supplementary Table 1**). Further, iRep values correlate as well as PTRs with growth rates determined from counts of colony forming units from the Korem *et al.* dataset (Fig. 2b).

**Figure 2.**
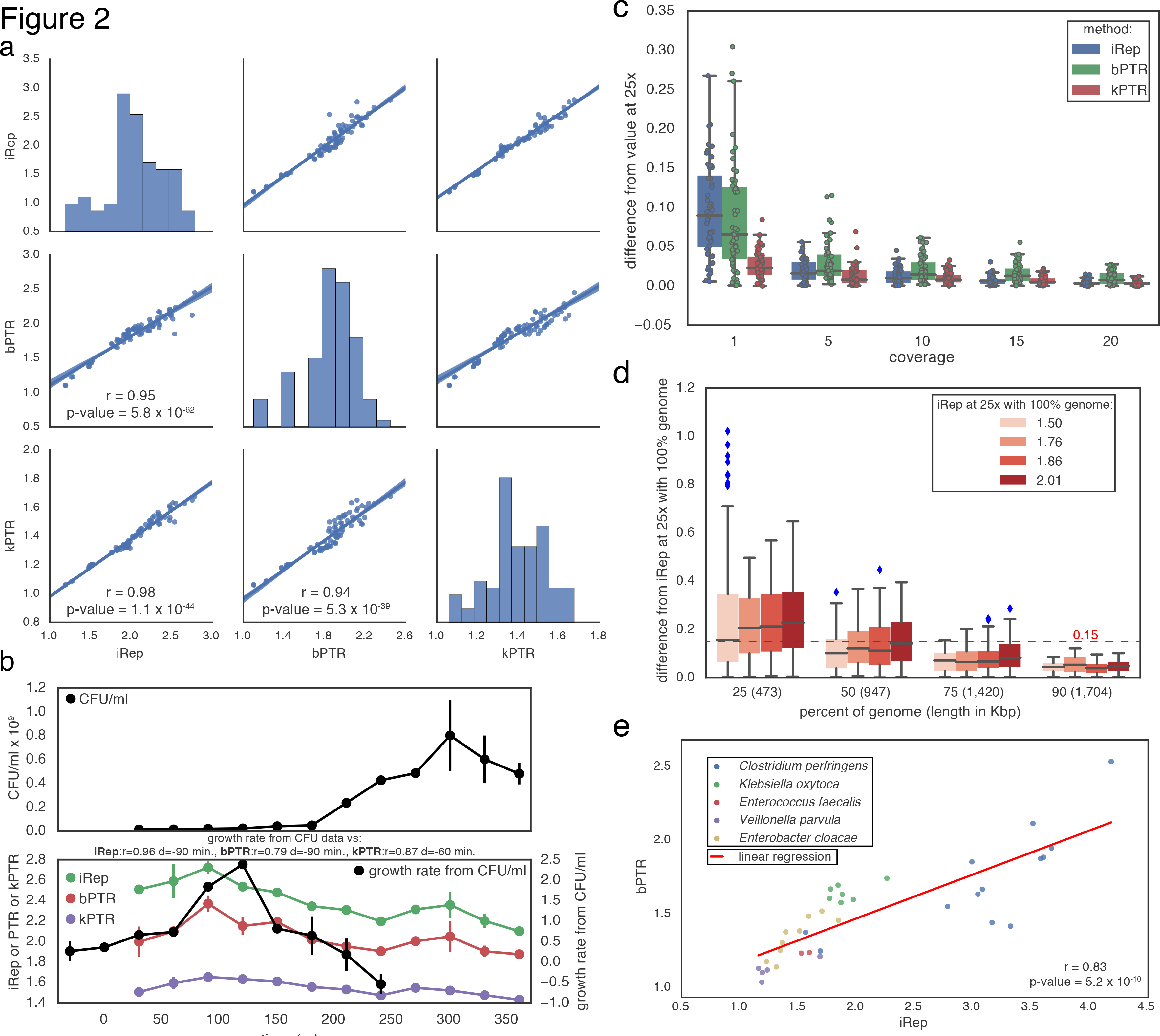
iRep can be used to measure *in situ* growth rates. (**a**) iRep, bPTR, and kPTR measurements made for cultured *Lactobacillus gasseri^6^* were compared (r = Pearson’s r value), showing strong agreement between all methods. (**b**) Colony forming unit (CFU) counts were available for a subset of these samples^6^, and used to calculate growth rates (n = 2). All methods were highly correlated with CFU-derived rates after first accounting for the delay between start of genome replication and observable change in population size (as noted previously^6^). Replication rates from CFU data were adjusted by-90 minutes before plotting, and by variable amounts for calculating correlations with sequencing-based rates (best correlation shown; d = time adjustment). (**c**) Using the *L. gasseri* data, minimum coverage requirements were determined for each method by first measuring the growth rate at 25x coverage, and then comparing to values calculated after simulating lower coverage. This shows that ≥5x coverage is required. (**d**) The minimum required genome fraction for iRep was determined by conducting 100 random fragmentations and subsets of the *L. gasseri* genome. Sequencing was subset to 5x coverage before calculating iRep to show the combined affect of low coverage and missing genomic information. With ≥75% of a genome sequence, most iRep measurements are accurate ±0.15. (**e**) iRep and bPTR measurements were calculated using five genome sequences assembled from premature infant metagenomes, showing that these methods are in agreement in the context of microbiome sequencing data.

We tested the minimum sequencing coverage required for iRep, kPTR and bPTR using data from culture-based *Lactobacillus gasseri* experiments from the Korem *et al.* study^6^. We first subsampled reads to achieve 25x coverage of the genome and then calculated replication rates to use as reference values. Then, the dataset was subsampled to lower coverage values and the replication rates re-calculated. Comparing these rates to the reference values enabled evaluation of the amount of noise introduced by increasingly lower coverage. Results show that all methods are affected by coverage and that, while kPTRs are the most precise at 1x coverage, all methods are reliable when the coverage is≥ 5× (Fig. 2c **and Supplementary Table 2**).

As a main goal of the iRep method is to obtain growth rate information when only draft quality genomes are available, we evaluated the minimum percentage of a genome required for obtaining accurate results by conducting a random genome subsampling experiment (**Fig. 2d, Supplementary Fig. 1, and Supplementary Table 3**). iRep values were determined for *L. gasseri* cells sampled when growing at different rates^6^, and then compared with values determined from genomes subset to various levels of completeness. Results indicate that ≥75% of the genome sequence is required for accurate measurements. As shown below, this standard can be met for a significant number of genomes recovered from metagenomic datasets.

The human microbiome includes organisms with genomes that are sufficiently similar to complete isolate genomes to enable ordering and orienting of draft genome fragments, making it possible to calculate both iRep and bPTR for comparative purposes. We conducted such an analysis using five genomes reconstructed in a previously published metagenomics study of premature infants^20^. Importantly, this strategy differs from kPTR in that the reads were mapped to the genome reconstructed from the infant gut samples to achieve more robust results than would be achieved using a public database-derived isolate genome. The correct ordering of the scaffolds in the reconstructed genome was confirmed based on both coverage patterns and cumulative GC skew (**Supplementary** Fig. 2). For 36 possible comparisons involving organisms with iRep values ranging between 1.2 and 4.2, there was a strong correlation between iRep and bPTR values (Fig. 2e). Given this, and the analysis of the Korem *et al.* experimental data described above, we conclude that iRep values are accurate measurements of replication rates.

Although a few reference genomes were similar enough to reconstructed genomes to facilitate scaffold ordering, the reference genomes were from organisms too distantly related to those being studied to enable accurate growth rate calculations. For the five genomes with available similar reference genomes (average nucleotide identity 91-99%), as much as 19.5% of reference genomes was not represented by metagenome reads (min. = 1.6%, average = 13.5%), compared with essentially perfect mapping to reconstructed genomes (**Supplementary Fig. 3 and Supplementary Table 4**).

Another opportunity to compare iRep and bPTR measures of replication rates was provided by an unusual case where a very large genome fragment was reconstructed from a highly complex groundwater metagenome. We manually curated the majority of a recently reported novel draft genome^21^ into a single scaffold (~2.5 Mbp). The scaffold contains both the origin and terminus of replication, as identified both by coverage and cumulative GC skew (Fig. 3). The bPTR value of 1.20 agrees with the iRep value of 1.23. Importantly, it would not have been possible to obtain this information based on mapping to complete reference genomes because this is the first sequence for an organism affiliated with a novel genus within the Deltaproteobacteria^21^. This finding demonstrates the iRep method in the context of a very complex natural environment.

**Figure 3.**
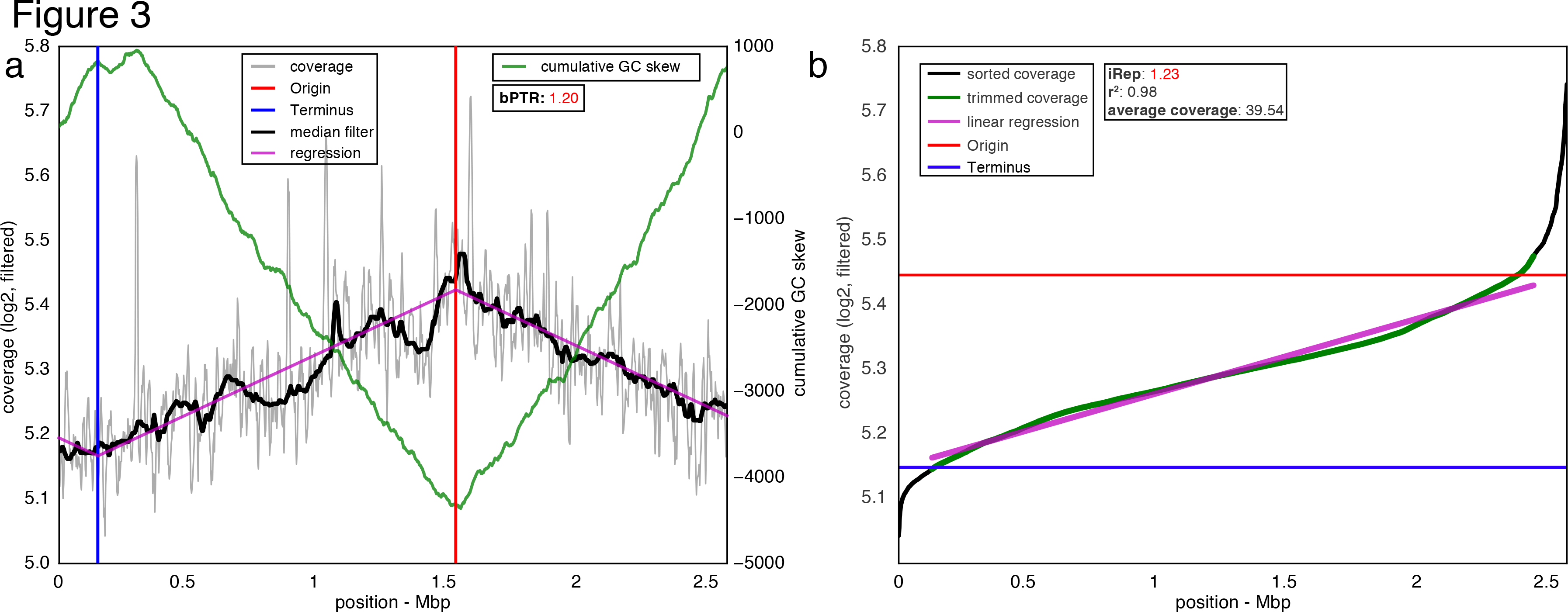
iRep and bPTR calculations agree for a novel Deltaproteobacterium sampled from groundwater^21^. (**a**) bPTR was calculated after determining the origin and terminus of replication based on regression to coverage calculated across the genome. Coverage was calculated for 10 Kbp windows sampled every 100 bp (**see Online Methods**). The ratio between the coverage at the origin and terminus was determined after applying a median filter. The cumulative GC skew pattern confirms the genome assembly and locations of the origin and terminus of replication. (**b**) iRep was determined by first calculating windows, as was done for bPTR, and then the resulting values were sorted. High and low coverage windows were removed, and then the slope of the remaining (trimmed) values was determined and used to evaluate the coverage at the origin and terminus of replication: iRep was calculated as the ratio of these values. (r^2^ was calculated between trimmed data and the linear regression).

### Replication rates from environmental and human-associated microbiomes

We obtained 275 iRep measurements using 202 genomes reconstructed as part of a study of premature human infant gut microbiomes^20^, and 51 genomes that we reconstructed from an adult human microbiome dataset^16^ (**Fig. 4a, Supplementary Tables 5-7, and Data Availability**). In infant microbiomes, members of the Firmicutes had the highest growth rates and Proteobacteria had the highest median growth rates (**Fig. 4b**). For 54 instances in which both iRep and kPTR could be determined, there was no strong correlation between the values (Pearson’s r = 0.45, **Supplementary Fig. 4, Supplementary Tables 5 and 7**). Because of the strong correlation between these methods when the organisms were represented by reference genomes (**Fig. 2a–b**), we attribute this disparity to a lack of appropriate complete genome sequences for calculating kPTR (**Supplementary Fig. 3**).

**Figure 4.**
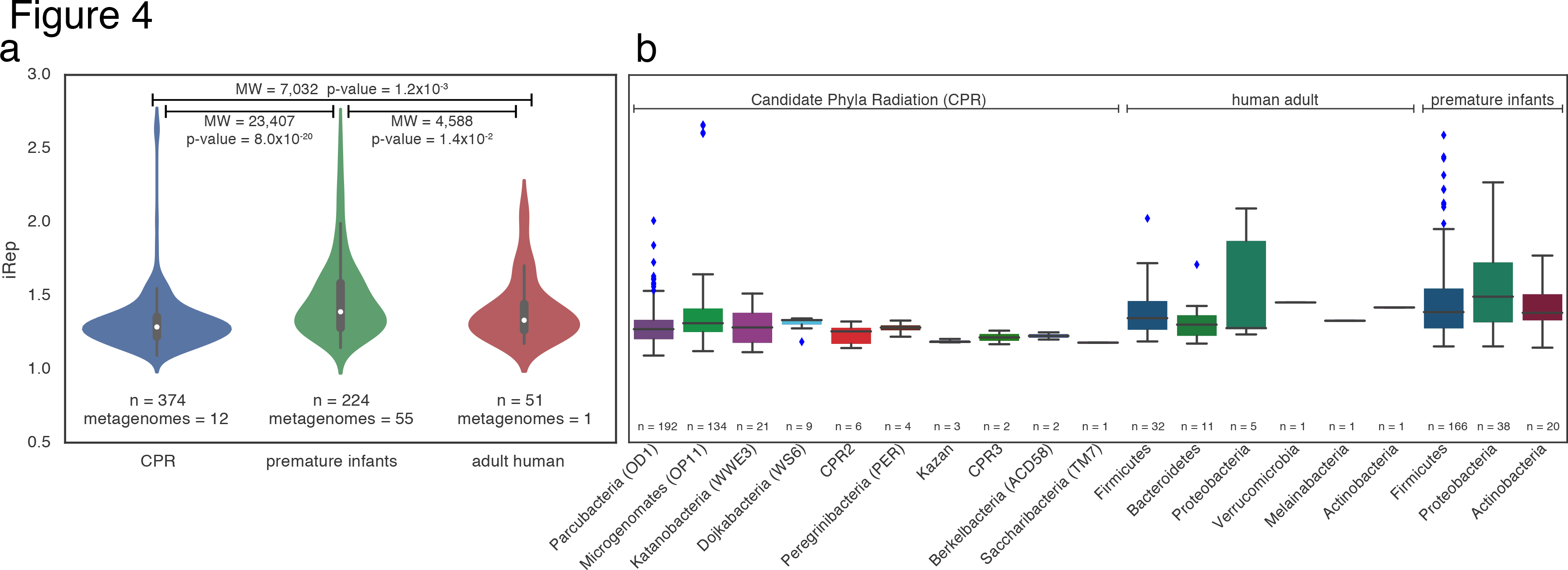
Growth rates for Candidate Phyla Radiation (CPR) and human microbiome-associated organisms were measured and compared across studies (**a**; MW = Mann-Whitney, n = number of measured growth rates), and compared based on taxonomic affiliation (**b**).

Using iRep, we obtained growth rates for 51 of the 54 genomes reconstructed from an adult human microbiome sample that were ≥75% complete (**see Methods; Fig. 4 and Supplementary Table 6**). Due to a lack of overlap with reference genomes, the kPTR method returned only three values, none of which were credible because all were <1 (**Supplementary Table 9**). Similarly, our attempt at selecting complete reference genomes for bPTR resulted in only five measurements (**Supplementary Fig. 5**). Even for these five cases, on average only 94% (min. = 88%, max. = 98%) of each complete reference genome was covered by metagenome sequences.

The Candidate Phyla Radiation (CPR) is a major subdivision within domain Bacteria known almost exclusively from genome sequencing^12^. Almost nothing is known about the growth rates of these enigmatic organisms. We measured 374 growth rates from CPR organisms using a time series of samples and 98 distinct genome sequences reconstructed from those datasets^12^ (**Supplementary Table 10**). Only 33 of the iRep values were calculated using complete genome sequences. One member of the CPR superphylum Microgenomates (OP11) exhibited some of the highest iRep values observed across CPR and human gut associated microorganisms (**Fig. 4b**). However, only 9.1% of iRep values from CPR organisms were >1.5, compared with 34.4% of premature infant and 17.6% of adult human microbiome measurements. Distributions of iRep values for organisms in premature infant and adult microbiomes were significantly different, but more similar to each other than to the CPR (**Fig. 4a**). The median iRep values for CPR bacteria are similar, but lower than those for human microbiome bacteria (CPR = 1.29, premature infant = 1.39, and adult = 1.33). Overall, the results demonstrate generally slower growth rates for CPR bacteria, as well as the applicability of the iRep method to bacteria in communities with very different levels of complexity.

### Microbial responses to antibiotics administered to premature infants

Samples were collected during periods of antibiotic administration for five of the ten studied infants^20^ (**Supplementary Fig. 6**). To measure microbial responses to antibiotics, we compared iRep values from samples collected within five days after antibiotic administration to values from other time points. This showed that organisms have significantly higher median growth rates after administration of antibiotics than at other times (**Fig. 5a**). Fast growing organisms were from the genera *Staphylococcus, Klebsiella, Lactobacillus, Escherichia*, and *Enterobacter* (iRep >1.8; **Supplementary Table 5**).

**Figure 5.**
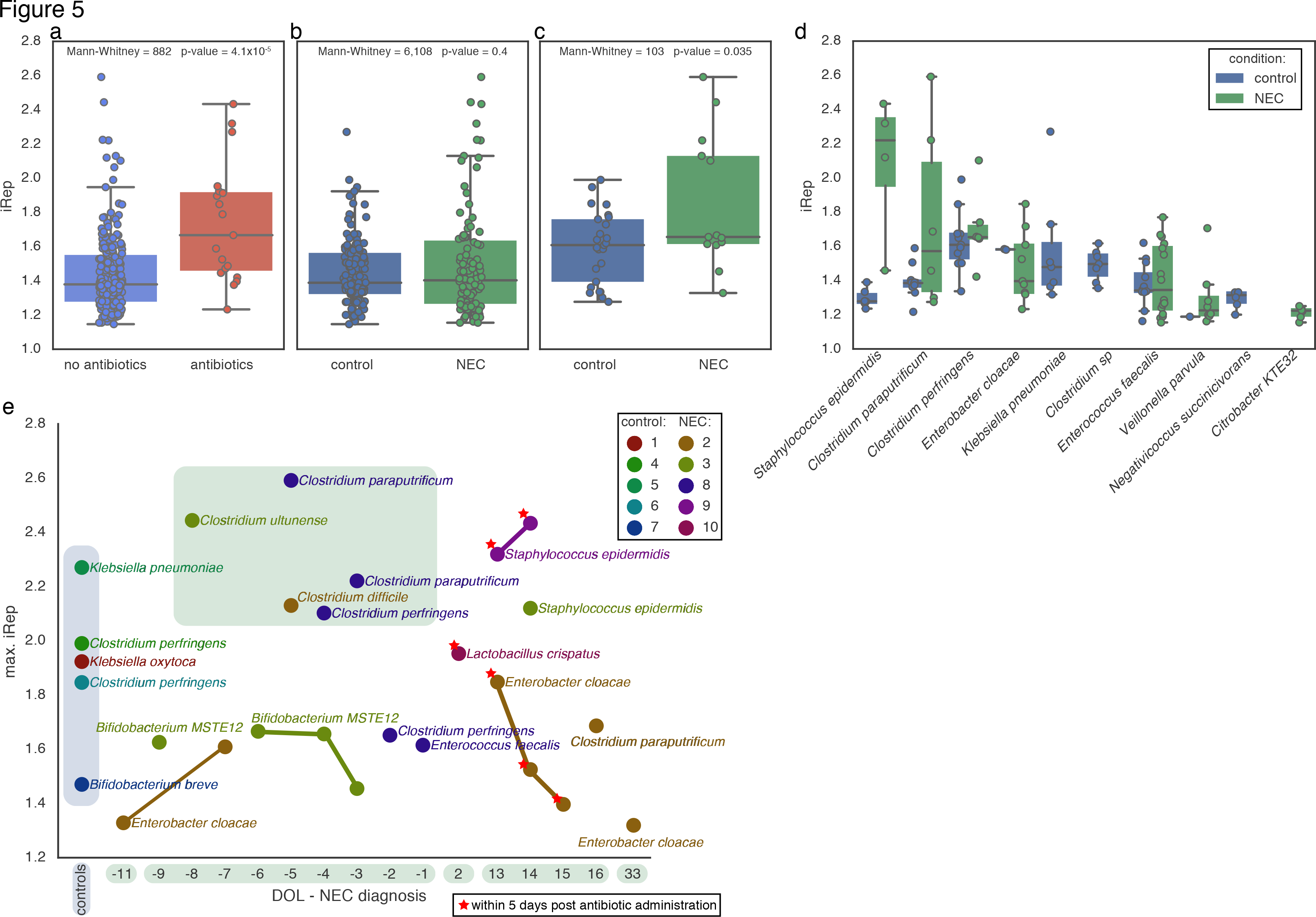
Elevated growth rates are associated with antibiotic administration and were detected prior to onset of necrotizing enterocolitis (NEC) in premature infants. iRep distributions were compared (**a**) between samples collected during or within five days after antibiotic administration and samples from other time points, (**b**) between samples collected from NEC and control infants, (**c**) between the highest iRep values from samples collected from NEC and control infants after excluding samples collected during or within five days of antibiotic administration. (**d**) Comparison of iRep values measured for different species sampled from NEC and control infants. (**e**) iRep for the fastest growing organism observed for each control infant (blue background), and for the fastest growing organism from each day of life (DOL) sampled for each NEC infant, reported relative to NEC diagnosis. High growth rates for members of the genus *Clostridium* were detected in infants surveyed prior to NEC diagnosis (green background).

### Fast replication rates for some bacteria in infants that developed necrotizing enterocolitis

The premature infant dataset consisted of 55 metagenomes collected from ten co-hospitalized premature infants, half of which developed necrotizing enterocolitis (NEC). While there is no statistically significant difference between iRep values from control and NEC infant microbiomes, it is striking that many of the highest values observed in the study were from NEC infants (Fig. 5b). After controlling for antibiotic administration and comparing only the highest iRep value observed in each sample, there is a modest, but statistically significant difference between iRep values from NEC and control infant microbiomes (Fig. 5c). *Staphylococcus epidermidis* growth rates were significantly higher in NEC versus control infants (Mann-Whitney p-value = 7.1x10^-3^). *Clostridium perfringens, Enterobacter cloacae*, and *Clostridium paraputrificum* all exhibited high growth rates in NEC infants, although some high growth rates for these organisms were also detected in control infants (**Fig. 5d**). Intriguingly, prior to development NEC, we detected high growth rates for four *Clostridum* species (**Fig. 5e**). Although replicating quickly in control infant microbiomes, *Klebsiella pneumonia* was detected infrequently in infants that developed NEC, and no iRep values could be determined.

### Microbial community dynamics measured by iRep and absolute abundance

Raveh-Sadka *et al.* measured absolute cell counts per gram of feces collected using droplet digital PCR (ddPCR) as part of the premature infant microbiome study^20^. Using these measurements we scaled species relative abundance values in order to track absolute changes in population sizes across samples (**Supplementary Table 5 and Supplementary Fig. 6**). For most of the infants in the study, iRep and both relative and absolute abundance values could be determined for most of the bacterial populations. Interestingly, despite fast growth rates observed for organisms from the genus *Clostridium* prior to NEC diagnosis, most of these organisms remained at low absolute abundance.

Typically, doubling times are calculated for organisms growing in pure culture without resource limitation or host suppression. We used the absolute abundance of *Klebsiella oxytoca* following antibiotic administration to calculate a realized doubling time of 19.7 hours (**Fig. 6a**). The iRep values for *K. oxytoca* during this period were consistently high (1.78-1.92), as required for the population growth that was well described by an exponential equation (r^2^ = 0.97). Notably, *K. oxytoca* was essentially the only organism present during this four-day period.

**Figure 6.**
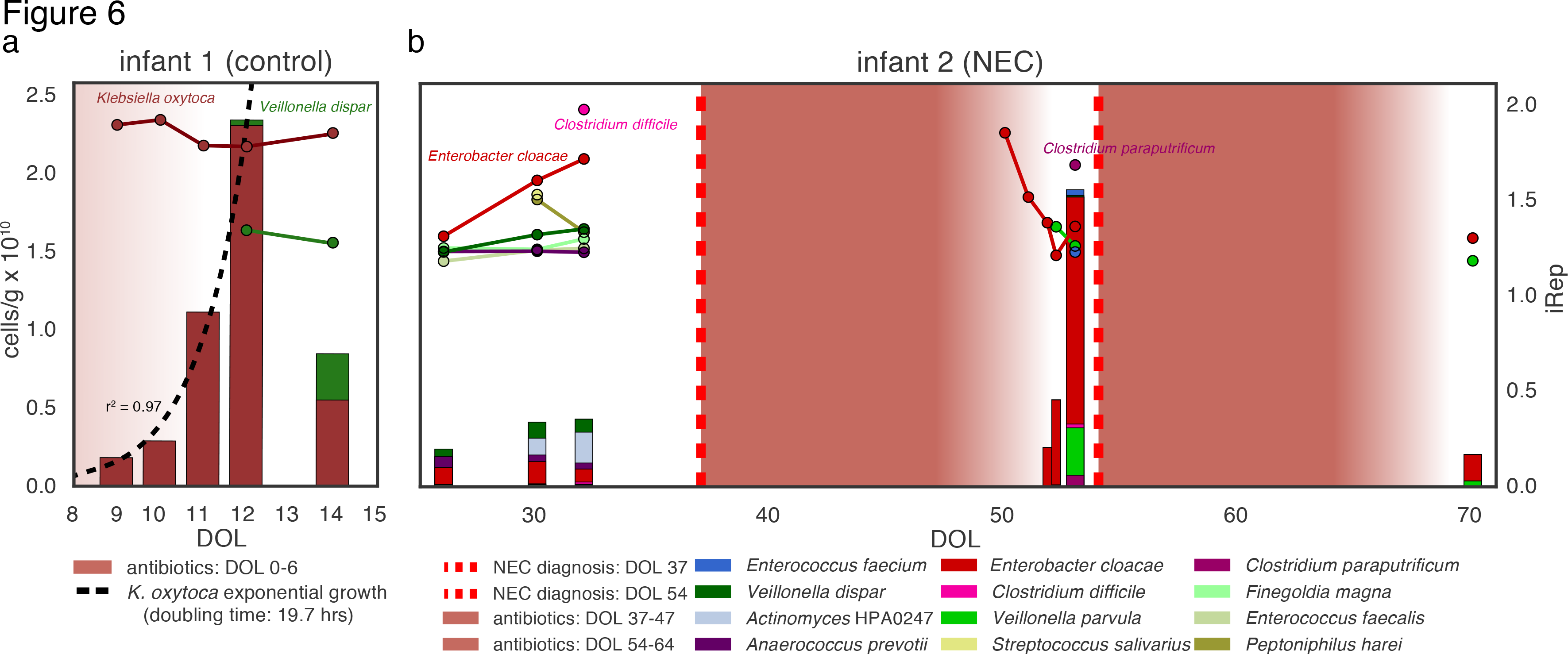
Absolute abundance (bars, left axis) and iRep (scatter plot, right axis) for bacteria associated with two premature infants (**a-b**; DOL = day of life). The five days following antibiotic administration are indicated using a color gradient. (**a**) Exponential growth was determined by regression to *K. oxytoca* absolute abundance values. (**b**) Infant 2 was diagnosed with two cases of necrotizing enterocolitis (NEC) during the study period.

In one infant, high iRep values were observed for *Clostridium difficile* and *Enterobacter cloacae* prior to the first NEC diagnosis, although these organisms remained at low absolute abundance (Fig. 6b). Total cell counts were low following antibiotic treatment; however, this period was associated with high *E. cloacae* growth rates and a subsequent 2.7-fold increase in population size prior to the second NEC diagnosis. Interestingly, low-abundance *Clostridium paraputrificum* was also growing quickly prior to the second diagnosis.

A clear finding from analysis of the growth rates for bacteria in multi-species consortia in the premature infant gut is the general lack of correlation between high iRep values and increased population size in the subsequently collected sample (**Supplementary Fig. 6**). It is important to note that at least one day separates sampling time points in almost all cases.

## Discussion

We developed a method for determining bacterial growth rates *in situ* that makes use of draft-quality genome sequences, which are now routinely generated in metagenomics studies. Bacterial growth rates derived from iRep using these genomes were generally more accurate than those obtained using genomes from reference databases. The combination of obtaining genomes from metagenomes and iRep measurements from the corresponding read data provides a comprehensive view of microbiome membership, metabolic potential, and *in situ* activity.

Despite the premature infant gut microbiome having relatively consistent community composition over time, iRep analyses indicate that brief flashes of rapid growth are common during the colonization period, possibly due to rapidly varying conditions in the infant gut. Even transitory levels of high activity, especially for potentially pathogenic organisms, could hold medical significance.

iRep measurements provide information about activity around the time of sampling. The approach can be used broadly to probe the responses of specific bacteria to environmental stimuli. However, periods of fast bacterial growth may not lead to increased population size because other processes exert controls on absolute abundances (e.g., predation and immune responses). In a few cases where community complexity was low, fast growth rates did predict an increase in absolute cell numbers in subsequent samples (**Fig. 6 and Supplementary Fig. 6**).

An important finding relates to the faster bacterial growth rates after antibiotic treatment, an observation that we attribute to high resource availability following elimination of antibiotic sensitive strains. Interestingly, rapid growth rates of several different but potentially pathogenic organisms from the genus *Clostridium*, including *C. difficile*, precede some NEC diagnoses, consistent with NEC being a multi-faceted disease. Further studies that include more samples and infants may establish a link between rapid cell division and NEC.

An interesting question relates to how quickly organisms proliferate in the premature infant gut compared to the adult gut environment. Measurements in such environments are very challenging using alternative approaches that make use of isotopes. These studies typically target specific organisms, and the measurements have only recently become somewhat tractable for the human microbiome^22^. Large-scale comparisons using PTR are not possible due to a lack of complete reference genomes. Using iRep, we found that bacteria from premature infant gut microbiomes had higher growth rates compared with those from a more complex adult gut consortium. If future studies establish this as a general phenomenon, it may reflect greater levels of competition for resources or other factors related to gut development in adults compared to premature infants.

Candidate Phyla Radiation (CPR) organisms have been detected in a wide range of environments^23^. Together, they make up considerably more than 15% of bacterial diversity ^12,24^yet they are known almost exclusively from genomic sampling ^12,15,25–30^ Based on having small cells and genomes with only a few tens of ribosomes, it was inferred that these organisms grow slowly ^23,31^. Our analysis of CPR organisms sampled across a range of geochemical gradients directly demonstrated their slow growth rates. However, the analysis also showed that some CPR bacteria grow rapidly under certain conditions (**Fig. 4**). Symbiosis has been inferred as a general life strategy for these organisms ^12,15,25–30^, and has been demonstrated in a few cases ^32–35^. Rapid growth of CPR bacteria may require rapid growth of host cells. If CPR cells typically depend on a specific bacterial host, as is the case for some Saccharibacteria (TM7)^34^, growth rate measurements may provide insights into possible host-symbiont relationships, paving the way for co-cultivation studies.

An important objective for microbial community studies is the establishment of models that can accurately predict microbial community dynamics and function under changing environmental conditions. Prior to the current study, these models could include growth rate information derived from laboratory experiments involving isolates, inferred from fixed genomic features such as 16S rRNA gene copy number or codon usage bias^36^, or from *in situ* measurements such as PTR^6^. We used iRep to quantify replication rates for most bacteria in infant gut microbial communities and found that the rates can be highly variable (**Fig. 5 and Supplementary Fig. 6**). Such measurements could be used in models that seek to understand microbial ecosystem functioning, allowing incorporation of organism-specific behavior throughout the study period. Importantly, the iRep methods can be applied to essentially all bacteria, regardless of how distantly related they are to previously studied species, to identify actively growing populations in any ecosystem, and to track organism responses to changing conditions. Such information has the potential to generate new understanding of relationships between bacteria and biogeochemical processes or health and disease outcomes.

## Online Methods

### Calculating bPTR for complete genomes

Our implementation of the PTR method (**see Code Availability**) differs from the method described by Korem *et al^6^* in several key respects. To distinguish between these two methods, we refer to our method as bPTR and the Korem *et al.* method as kPTR. Both methods involve mapping DNA sequencing reads to complete (or near-complete, in the case of bPTR) genome sequences in order to measure differences in sequencing coverage at the origin and terminus of replication. kPTR makes use of a database of reference genome sequences, whereas bPTR is designed to be more flexible and can use mapping of reads to any genome sequence. For our bPTR analyses, we used Bowtie2^37^ with default parameters for read mapping.

Both bPTR and kPTR can determine the location of the origin and terminus of replication of growing cells by identifying coverage “peaks” and “troughs” associated with these positions. Identification of the origin and terminus of replication requires measuring changes in coverage along the genome sequence. This is accomplished by calculating the average coverage over 10 Kbp windows at positions along the genome separated by 100 bp. To increase the accuracy of results, a mapping quality threshold can be used in which both reads in a set of paired reads are required to map to the genome sequence with no more than a specified number of mismatches (this option is unique to bPTR). Since highly conserved regions, strain variation, or integrated phage can result in highly variable coverage, high and low coverage windows are filtered out of the analysis. Coverage windows are excluded if the values differ from the median by a factor greater than 8 (threshold also used by kPTR), or if the values differ from the average of 1,000 neighboring coverage windows by a factor greater than 1.5 (threshold unique to bPTR). If more than 40% of the windows are excluded, no bPTR value will be calculated (threshold also used by kPTR). The origin and terminus are identified by fitting a piecewise linear function to the filtered, log2-transformed coverage values. Coverage values are log2-transformed to improve fitting, but the transformation is reversed prior to calculating bPTR. Fitting is conducted as described by Korem *et al.* by non-linear least squares minimization using the Levenberg-Marquardt algorithm implemented by lmfit ^38^.

Piecewise linear function modified from Korem *etal^6^:*

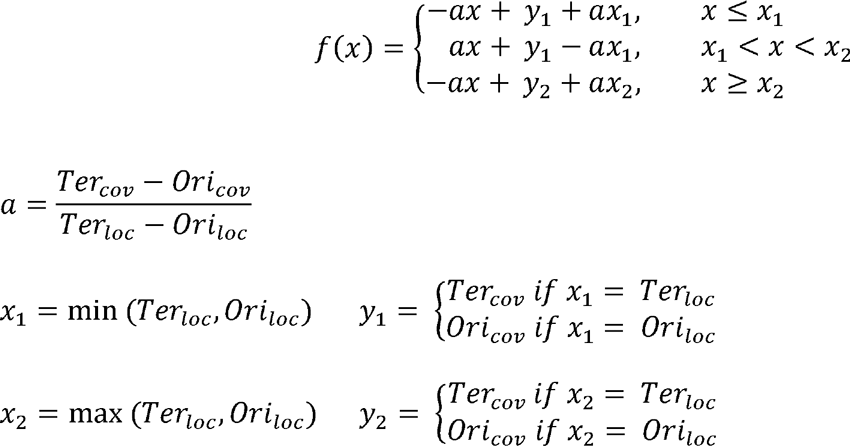

*Ori_loc_* and *Ter_loc_* refer to the locations of the origin and terminus of replication, respectively, and *Ori_cov_* and *Ter_cov_* refer to log_2_-transformed coverage at those positions. All *x* values refer to positions on the genome, and *y* values to log_2_-transformed coverage values. The fitting is constrained such that *Ori_loc_* and *Ter_loc_* are separated by 45-55% of the genome length^6^. In order to reduce the amount of noise introduced by fluctuations in sequencing coverage, a median filter is applied to the coverage data before calculating bPTR. This smoothing operation replaces the coverage value at each position with the median of values sampled from the 1,000 neighboring windows. The log_2_-transformed, median-filtered values corresponding with *Ori_loc_* and *Ter_loc_* (*Ori_cov-med_* and *Ter_cov-med_*, respectively) are used to calculate bPTR.

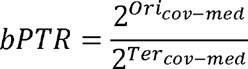

*Ori_loc_* and *Ter_loc_* are determined based on sequencing from each available sample. In order to calculate bPTR using the same positions for all samples, consensus *Ori_loc_* and *Ter_loc_* positions are determined by finding the circular median of the positions determined from each individual sample (all *Ori_loc_* and *Ter_loc_* positions with bPTRs ≥1.1 are considered), as is done for kPTR^6^. Once these values are determined, all bPTR values are re-calculated using the coverage at the consensus positions. It is important to note that *Ori_loc_* and *Ter_loc_* may vary depending on what samples are analyzed, and that this can be avoided by using GC skew to identify *Ori_loc_* and *Ter_loc_* (**see below**).

For bPTR we added the option to find *Ori_loc_* and *Ter_loc_* based on GC skew. GC skew is calculated over 1 Kbp windows at positions along the genome separated by 10 bp. Since *Ori_loc_* and *Ter_loc_* coincide with a transition in the sign (+/−) of GC skew, these positions can be identified as the transition point in a plot of the cumulative GC skew^39^ (for examples see **Fig. 3, Supplementary Fig. 2, and Supplementary Fig. 5**). These transition points are identified by finding extreme values in the cumulative GC skew data separated by 45-55% of the genome length. Once *Ori_loc_* and *Terloc* are identified, bPTR is calculated from median-filtered log_2_-transformed coverage values calculated over sliding windows as described above. bPTR provides visual representation of both coverage and GC skew patterns across genome sequences that enable verification of genome assemblies and predicted *Ori_loc_* and *Ter*_loc_positions (this visualization is not provided by kPTR).

### Calculating the Index of Replication (iRep) for complete and draft-quality genomes

iRep (**see Code Availability**) analyses are conducted by first mapping DNA sequencing reads to genome sequences with Bowtie2 (default parameters). Then, coverage is evaluated by calculating the average coverage over 10 Kbp windows at positions along the genome separated by 100 bp. For genomes in multiple pieces, the fragments are combined in an arbitrary order (**see below**) before performing sliding window calculations. To increase the accuracy of results, a mapping quality threshold can be used in which both reads in a set of paired reads are required to map to the genome sequence with no more than a specified number of mismatches. Coverage windows are filtered out of the analysis if the values differ from the median by a factor greater than 8. Coverage values are log_2_-transformed and then sorted from lowest to highest coverage. Because the coverage windows are re-ordered in this step, it does not matter if the correct order of genome fragments is unknown. The lowest and highest 5% of sequences are excluded, and then the slope of the remaining coverage values is determined by linear regression. As with bPTR, log_2_-transformations are conducted to improve regression analysis, but are removed before comparing coverage values. iRep, which is a measure of the ratio between *Ori_cov_* and *Ter_CO_y*, can be determined based on the slope (m) and total length of the genome sequence *(l)*:

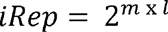

Since partial genome sequences will include a random assortment of genome fragments, the coverage trend determined from the available sequence will be representative of the coverage trend across the complete genome. Because of this, only the slope of coverage values across the partial genome, and the length of the partial genome need to be known (the length of the complete genome sequence is not needed).

Several quality thresholds are used to ensure the accuracy of iRep measurements: i) coverage depth must be ≤5x, ii) ≤98% of the genome sequence must be included after filtering coverage windows, and iii) r^2^ values calculated between the coverage trend and the linear regression must be ≥0.90. These criteria are important because they ensure that enough sequencing data is present to achieve accurate measurements, and that the genome sequence is appropriate for the analysis. The 98% genome sequence coverage threshold differs from the genome completeness requirement in that this is not a measure of the quality of the genome assembly, but rather a measure of the overlap between a genome sequence and the sequencing data. Low values would indicate that the genome used for mapping is not appropriately matched with an organism present in the system. Likewise, having a strong fit of the linear regression to the coverage data indicates that sequencing coverage calculations are not influenced by strain variation, choice of an inappropriate genome sequence, or other factors that may skew growth rate measurements.

Both PTR methods involve calculations based on only two data points *(Ori_cov_* and *Ter_cov_).* In contrast, iRep uses coverage trends determined across an entire genome sequence, and thus is less susceptible to noise in sequencing coverage or errors in the prediction of *Ori_loc_* and *Ter_loc_*. Further, since both PTR methods involve predicting *Ori_loc_* and *Ter_loc_* based on data from multiple samples, the same positions may not be chosen for different analyses. This makes it difficult to reproduce and compare results (an issue that can be avoided by predicting *Ori_loc_* and *Ter_loc_* using cumulative GC skew and bPTR). iRep calculations do not depend on analysis of multiple samples, and thus results will not change based on what samples are included in an analysis. Since the order of genome fragments need not be known when calculating iRep, the method is not affected by genome assembly errors, which are present even in some genome sequences reported to be complete (**Supplementary Fig. 5**).

### Determining the minimum coverage required for iRep analysis

*Lactobacillus gasseri* data from the Korem *et al.^6^* study was used to determine the minimum coverage required for iRep, bPTR, and kPTR. Reads from each sample were first mapped to the complete genome sequence, and then sub sampled to 25x before calculating iRep, bPTR, and kPTR. Then, each mapping was further subsampled to lower coverage levels (20x, 15x, 10x, 5x, and 1x) and growth rates were re-calculated using each method. Comparison of these values to those determined at 25x coverage enable quantification of the amount of noise introduced by increasingly lower coverage (**Fig. 2c and Supplementary Table 2**).

### Determining the minimum genome fraction required for iRep analysis

The *L. gasseri* data from Korem *et al ^6^* sub sampled to 25x coverage was also used to test the minimum fraction of a genome required for obtaining accurate iRep measurements. Four samples representing iRep values between 1.50 and 2.01 were selected in order to test the effect of missing genomic information across a range of replication rates. Genome subsampling experiments were conducted on each sample in order to evaluate the amount of noise introduced by missing genomic information. For each tested genome fraction (90%, 75%, 50%, and 25%), iRep was calculated for 100 random genome subsamples. For each subsample, the genome was fragmented into pieces with lengths determined by selecting from a gamma distribution modeled after the size of genome fragments expected for draft-quality genome sequences (**Supplementary Fig. 1**). Once fragmented, the pieces were randomly sampled until the desired genome fraction was achieved. Partial fragments were included in order to prevent the desired genome fraction size from being exceeded. In order to ensure that the results were accurate even when sequencing coverage is low, iRep calculations were conducted after subsampling reads to 5x coverage. iRep values calculated after subsampling were compared to values determined at 25x coverage with the complete genome sequence in order to measure the combined affect of lower coverage and missing genome sequence information (**Fig. 2d and Supplementary Table 3**).

### Comparative analyses of replication rate methods

iRep, bPTR, and kPTR were calculated for all samples from the Korem *et al.*^6^ study (**Supplementary Table 1**). However, only the *L. gasseri* experiments were sequenced to a high enough depth to enable comparison with iRep (**Fig. 2a**). For a subset of these data, growth rates could also be calculated based on counts of colony forming units (CFU/ml)^6^ (**Fig. 2b**). Pearson’s correlations were calculated between growth rates based on CFU/ml data and iRep, bPTR, and kPTR, after first accounting for the time delay between start of genome replication and observable change in population size (as previously noted^6^). The time delay was determined independently for each method as the delay that resulted in the highest correlation.

iRep and bPTR values were compared for a novel Deltaproteobacterium after manually curating the draft genome sequence recently reported by Sharon *et al^21^* (see below). Reads from the GWC2 sample from Brown *et al.^12^* were used to conduct the analysis (**Fig. 3**). For this comparison, and all subsequent iRep and bPTR calculations, coverage was calculated based on reads that mapped to the genome fragment with no more than two mismatches (**see above for details**). Although enough of the genome sequence was assembled in order to calculate bPTR, the results could not be compared with kPTR because a complete genome sequence was not available.

In order to further compare iRep and bPTR in the context of microbial community sequencing data, bPTR values were calculated using genomes reconstructed from the premature infant dataset^20^ that were ordered and oriented based on complete reference genome sequences (see below; Fig. 2e and Supplementary Table 4). Although these genomes were similar enough to reference genomes to facilitate ordering and orienting the sequences, the reference genomes themselves were too divergent to facilitate growth rate calculations (**see Results; Supplementary Fig. 1**), which prevented inclusion of kPTR in this analysis.

### Manual curation of novel Deltaproteobacterium genome

The genome sequence of a previously reported novel Deltaproteobacterium was manually curated. Unplaced or misplaced paired-read sequences were used to fill scaffolding gaps, correct local assembly errors, and extend scaffolds. Overlapping scaffolds were combined when the join was supported by paired read placements. The final assembled sequence was visualized to confirm that all errors had been corrected.

### Ordering and orienting draft genomes based on complete reference genomes

Reference genomes similar to draft genomes were obtained from NCBI GenBank. Genomes with aberrant GC skew patterns were not used for ordering draft genomes as they likely contain assembly errors. The average nucleotide identities (ANI) between each draft genome and associated reference genomes were calculated using the ANIm method^40^, and the reference genome with the highest ANI was chosen. Draft genome fragments were aligned to the reference genome using BLAST^41^, and any fragment with less than 20% alignment coverage was discarded. The remaining sequence was then aligned to the reference genome using progressive Mauve ^42^, resulting in an ordered and oriented genome to be used for calculating bPTR. These genomes were manually inspected and curated based on cumulative GC skew and genome coverage patterns based on graphs generated by the bPTR script (**Supplementary Fig. 2**).

### iRep measurements for premature infant metagenomes

Previously reconstructed genomes from the premature infant gut microbiome study^20^ were included in the iRep analysis if they were estimated to be ≥75% complete based on analysis of universal single copy genes (with no more than two duplicate genes). In order to maximize the number of iRep values that could be determined, custom read mapping databases were used for each metagenome. Each database was constructed by first including genomes reconstructed from the metagenome, and then by adding additional genomes reconstructed from other metagenomes from the same infant. This prioritizes genomes reconstructed from the metagenome used for mapping, but also attempts to include genomes from organisms that may have been present, but for which a genome sequence was not assembled.

Overlap in community membership across time-series studies results in the same genome sequence being reconstructed in multiple samples. Including highly similar or identical genome sequences in databases used for read mapping would lead to aberrant coverage calculations. This becomes a concern when including genomes reconstructed from additional samples in read mapping databases for iRep calculations. To prevent adding highly similar genomes to the databases, only the representatives of 98% ANI genome clusters (**see below**) were added to mapping databases, and only if a representative of the cluster was not already included. Consistent with clustering genomes based on sharing 98% ANI, iRep calculations were conducted based on coverage calculations determined from reads mapping to genomes with no more than two mismatches (**see above for details; Supplementary Table 5**).

### Clustering genomes based on average nucleotide identity (ANI)

Average nucleotide identity was determined between all pairs of genome sequences using the Mash algorithm^43^ (kmer set to 21). Clusters were defined by selecting groups of genomes connected by ≥98% ANI. Representatives of each cluster were chosen by selecting the longest genome with the largest number of single copy genes, and the fewest number of single copy gene duplicates.

### Comparison of iRep and kPTR measurements for premature infant gut metagenomes

The kPTR software from Korem *et al^6^* was run on the premature infant metagenomes^20^ (**Supplementary Table 8**). Comparisons between iRep and kPTR were made when it was possible to link the name of the genome provided by kPTR with the taxonomy given to reconstructed genome sequences (**Supplementary Table 5**).

### Genome binning and iRep measurements for adult human metagenomes

Genomes were binned from the adult human metagenome^16^ based on coverage, GC content, and taxonomic affiliation using ggKbase tools (ggkbase.berkeley.edu), as previously described^12,20^. Genome completeness was evaluated based on the fraction of universal single copy genes^20,44^ that could be identified (**Supplementary Table 6**). Genomes estimated to be ≥75% complete, with no more than two additional single copy genes, were used in the analysis. iRep was conducted using read mapping to genomes with ≤2 mismatches (**Supplementary Table 7**).

### bPTR and kPTR measurements from the adult human metagenome

The kPTR software from Korem *et al.^6^* was run on the adult human metagenome^16^ (**Supplementary Table 9**). bPTR calculations were conducted based on mapping metagenome reads to selected complete reference genomes (≤2 mismatches; **Supplementary Fig. 5**). Reference genomes for bPTR analysis were selected by searching scaffolds from reconstructed genome sequences against complete genomes from NCBI GenBank. The complete genome with the best BLAST hit to each reconstructed genome was selected for bPTR analysis.

### iRep measurements for Candidate Phyla Radiation (CPR) organisms

CPR genomes identified by Brown *et al.*^12^ to be ≥75% complete, with no more than two additional single copy genes, were selected for iRep analysis. These genomes were reconstructed previously from multiple metagenomes spanning a time-series field experiment. Reads from each of 12 metagenomes sequenced from groundwater filtrates, collected from serial 0.2 and 0.1 filters at six time points, were mapped to the genome sequences for iRep calculations (≤ 2 mismatches; **Supplementar Table 10**).

### Absolute abundance and doubling time determinations

Raveh-Sadka *et al.* determined the concentration of cells in each collected fecal sample using droplet-digital PCR^20^. In the current study, the population size of each species was determined by multiplying total cell counts by the fractional (relative) abundance calculated based on genome sequencing (**Supplementary Table 5**). These values were used to calculate realized doubling time for *Klebsiella oxytoca* (**Fig. 6**).

## Acknowledgements

Funding was provided by NIH grant 5R01AI092531, Sloan Foundation grant APSF-2012-10-05, and by the US Department of Energy (DOE), Office of Science, Office of Biological and Environmental Research under award number DE-AC02-05CH11231 (Sustainable Systems Scientific Focus Area and DOE-JGI) and award number DE-SC0004918 (Systems Biology Knowledge Base Focus Area). We thank T. Raveh-Sadka, B., Brooks, and D. Burstein for helpful discussions.

## Data and Code Availability

DNA sequencing reads are available from the NCBI Sequence Read Archive for the groundwater^12^ (SRP050083), premature human infant^20^ (SRP052967), and adult human^16^ (SRR3496379) microbiome projects. Genomes analyzed as part of this study are available from ggKbase for the groundwater^12^ <u>(ggkbase.berkeley.edu/CPR-complete-draft/organisms</u>), premature human infant <u>(ggkbase.berkeley.edu/project groups/necevent samples)</u>, and adult human <u>(ggkbase.berkeley.edu/LEY3/organisms</u>) datasets, as well as for the curated novel Deltaproteobacterium <u>(ggkbase.berkeley.edu/novel delta irep/organisms)</u>. CPR genomes are available from NCBI GenBank BioProject PRJNA273161. iRep and bPTR software are maintained under <u>https://github.com/christophertbrown/iRep</u> (v0.1-beta used in this analysis: <u>https://github.com/christophertbrown/iRep/releases/tag/v0.1-beta</u>).

## Supplementry Figures

**Supplementary figure 1.**
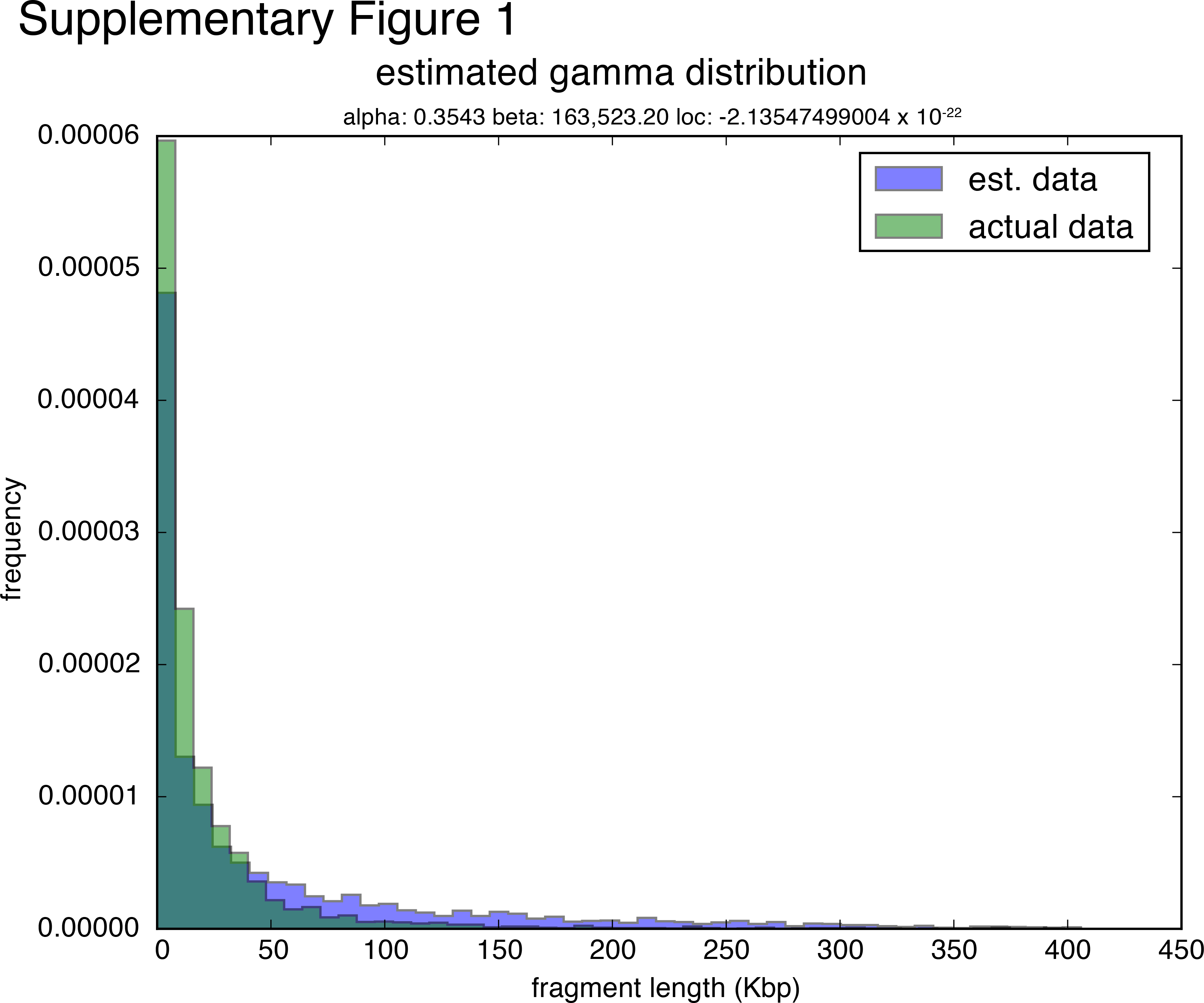
Gamma distribution used to simulate genome fragmentation for the genome completeness analysis (**Fig. 2d; see Online Methods**). Frequency of genome fragment sizes from the CPR genome dataset are compared with genome fragment sizes randomly sampled from a gamma distribution with parameters: alpha = 0.35, beta = 164,000. These parameters were first estimated by fitting to the CPR data, and then manually adjusted. Similarity between the two distributions shows that this gamma distribution can be used to approximate the level of genome fragmentation expected for draft-quality genome sequences.

**Supplementary figure 2.**
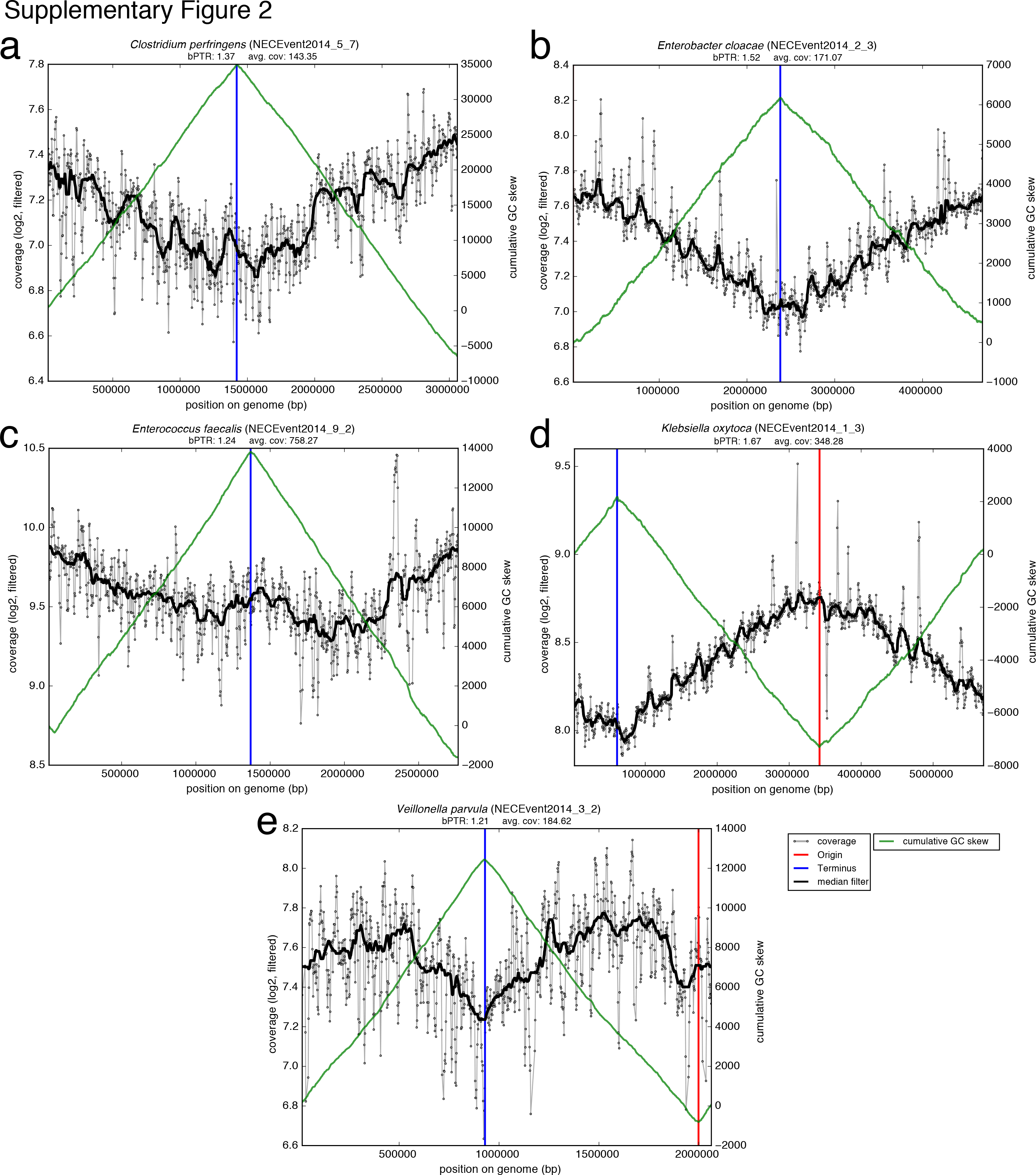
Coverage, GC skew patterns, and bPTR measurements for reconstructed genomes oriented and ordered based on complete reference genome sequences. (**a-e**) Read mapping was conducted using sequences from the sample used for genome recovery. bPTR was calculated after determining the origin and terminus of replication based on cumulative GC skew. Coverage was calculated for 10 Kbp windows calculated every 100 bp (extremely low and high coverage windows were filtered out; **see Online Methods**). bPTR was calculated as the ratio between the coverage at the origin and terminus after applying a median filter. Cumulative GC skew and coverage patterns confirm the ordering of genome fragments.

**Supplementary figure 3.**
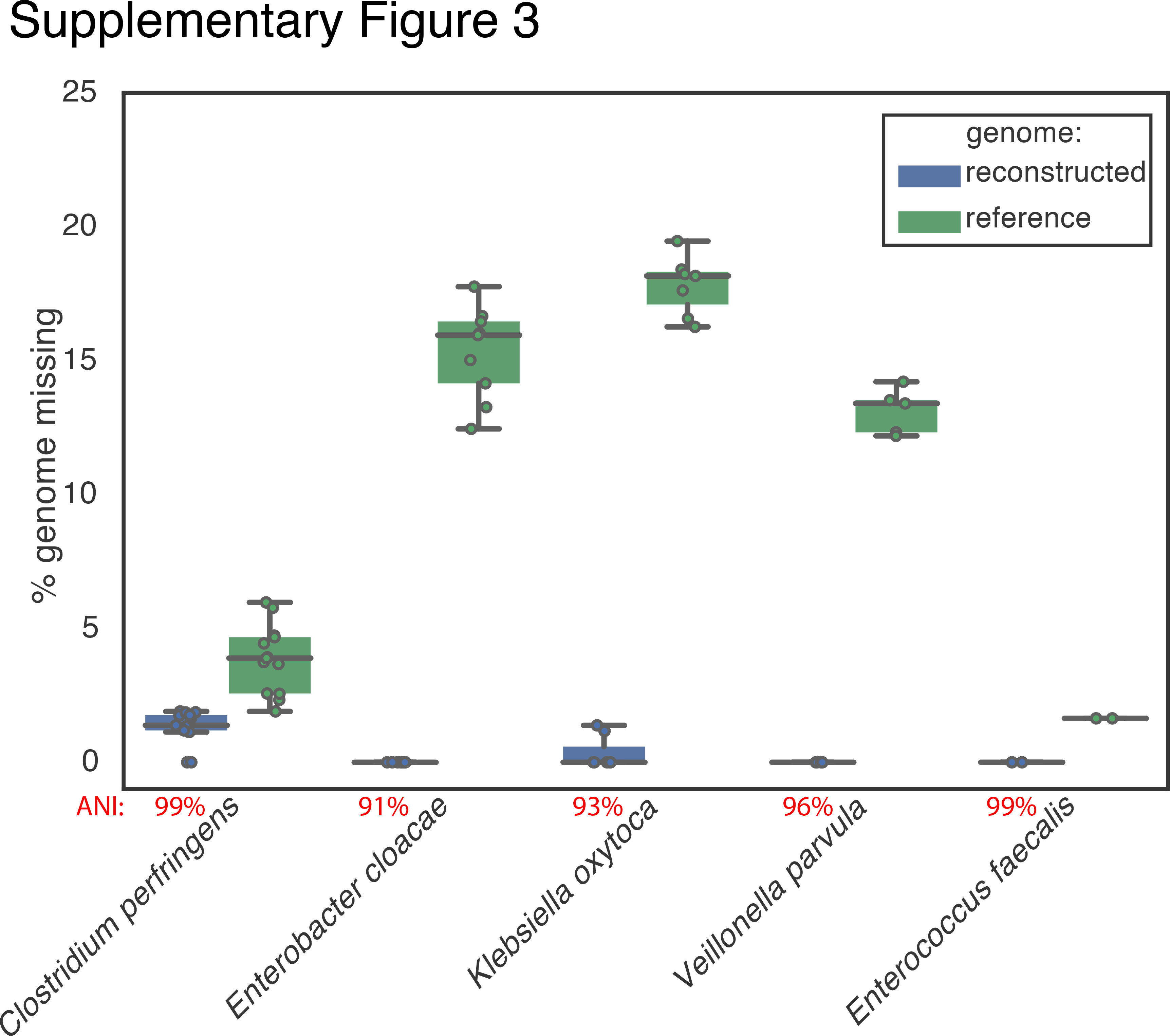
Reference genomes are not representative of organisms surveyed in the premature infant microbiome study. Reads were mapped to both reconstructed genomes and closely related reference genomes (**Supplementary Table 4**), and the percent of each genome covered by sequencing reads is reported. Average nucleotide identity (ANI) is reported between each reconstructed genome and the paired reference genome. The large fractions of reference genomes not represented by metagenome sequencing show that extensive genomic variation is present between surveyed and reference genomes, despite high ANI values in some cases.

**Supplementary figure 4.**
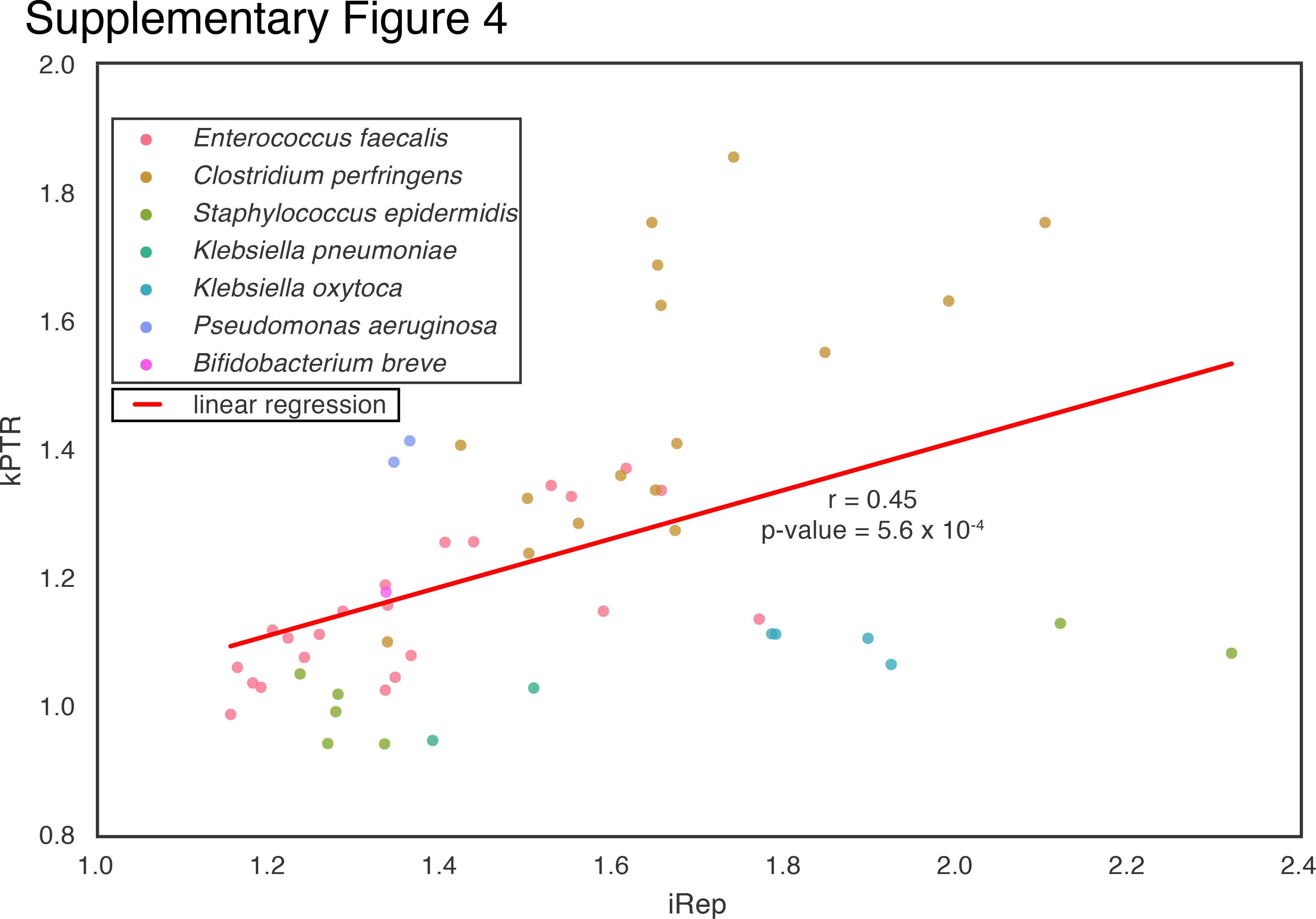
Growth rates determined by iRep (using reconstructed genomes) and kPTR (using reference genomes) are not in strong agreement for the premature infant study (r = Pearson’s r value).

**Supplementary figure 5.**
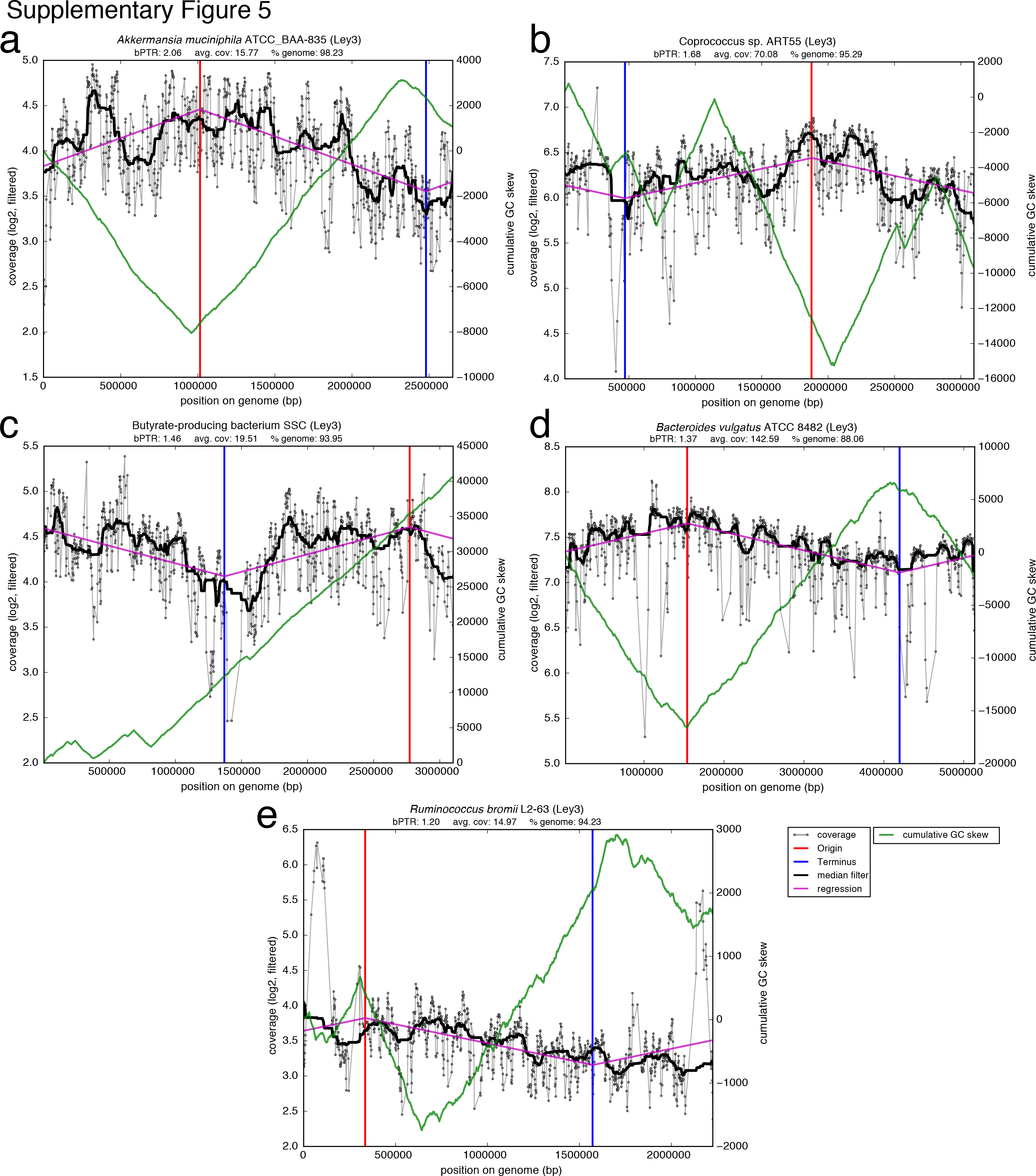
Coverage, cumulative GC skew patterns, and bPTR measurements for complete reference genomes with similarity to genomes from the adult human microbiome sample. (**a-e**) Reads from the adult human microbiome were mapped to complete reference genome sequences. Coverage was calculated for 10 Kbp windows every 100 bp (extremely low and high coverage windows were filtered out; **see Online Methods**). The origin and terminus of replication were determined based on coverage. bPTR was calculated as the ratio between the coverage at the origin and terminus after applying a median filter. Cumulative GC skew and coverage patterns suggest the presence of genomic variation or assembly errors for some genomes (**b-c, e**).

**Supplementary figure 6.**
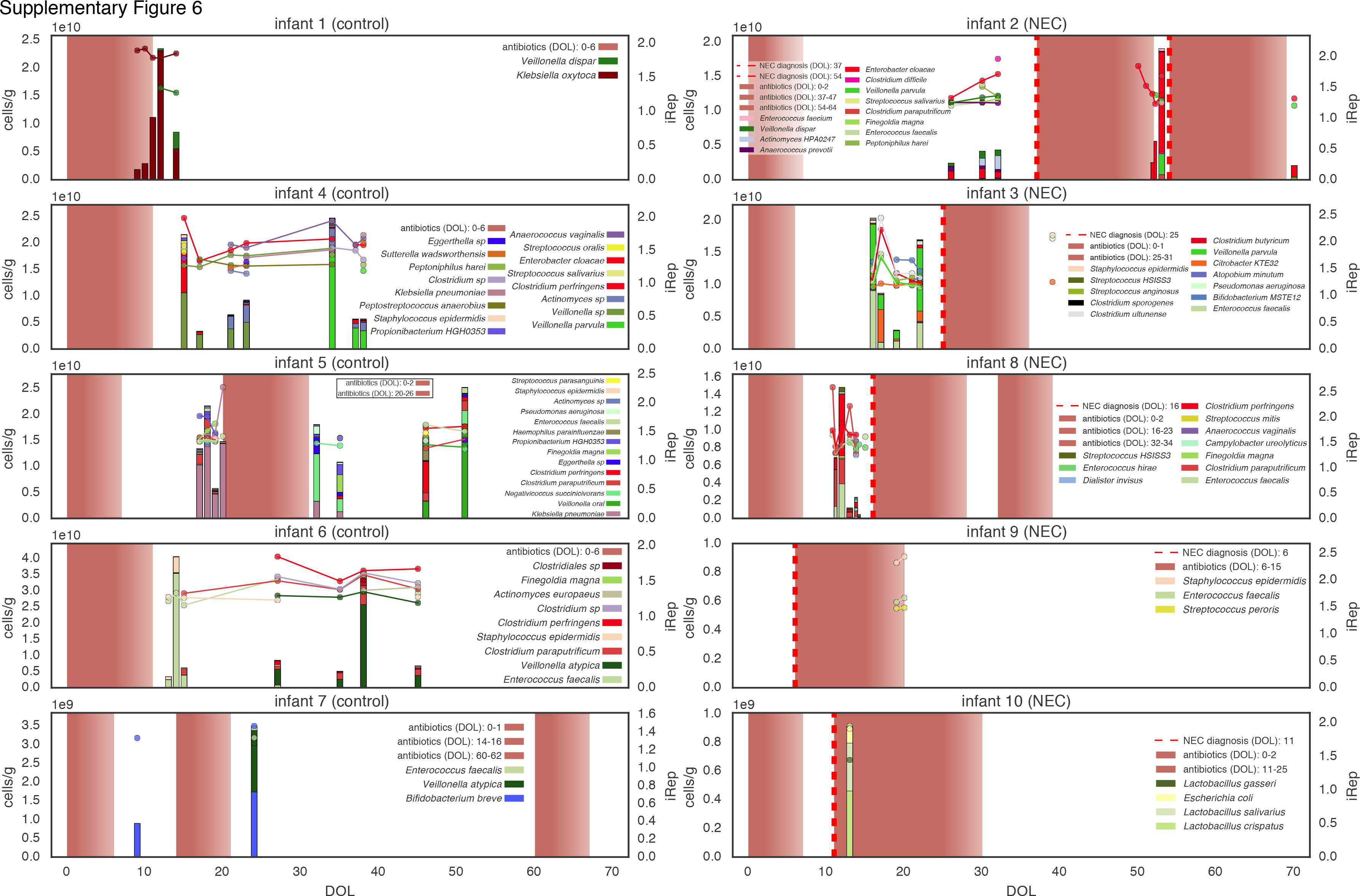
Absolute abundance (bars, left axis) and iRep (scatter plot, right axis) for bacteria associated with premature infants (DOL = day of life). The five days following antibiotic administration are indicated using a color gradient.

**Supplementary Table 1** iRep, bPTR, and kPTR measurements from Korem *et al.^6^* datasets. r^^^2 is the r^2^ value calculated between the sequencing coverage trend and regression used for calculating iRep, and % windows refers to the percent of coverage windows that passed the iRep filters. Coverage is the average sequencing depth calculated across the genome sequence.

**Supplementary Table 2** iRep, bPTR, and kPTR measurements for genome sequencing coverage analysis conducted using data from previously published experiments^6^. Target coverage is the level of coverage achieved after sub-sampling sequencing reads.

**Supplementary Table 3** Results of the analysis of the impact of genome coverage on growth rate determination using sequencing data from *Lactobacillus gasseri* culture experiments^6^. iRep was first calculated for the complete genome at 25x coverage, and then after sub sampling the genome 100 times for each targeted percent of the genome sequence. The analysis was repeated after simulating 5x coverage for the subsampled genome (target coverage).

**Supplementary Table 4** iRep and bPTR measurements calculated after ordering and orienting genomes reconstructed for organisms sampled as part of the premature infant dataset^20^. Values were calculated for all pairs of genomes and samples by mapping reads from the sample to the genome sequence (matched is whether or not the genome and sample were collected from the same infant). The analysis was conducted using both reconstructed and reference genome sequences (see ref. genome columns for reference genome data).

**Supplementary Table 5** iRep measurements for organisms associated with premature infant microbiomes^20^ (DOL = day of life; NEC = necrotizing enterocolitis). Absolute abundance (cells/g) was determined for each organism based on relative abundance and previously published ddPCR measurements of total cells/g of feces. The antibiotics column indicates whether or not antibiotics were administered at, or within five days prior to, the time of sample collection. The DOL-sample column indicates whether additional samples were collected on a particular day, the DOL (NEC) columns includes day of life relative to NEC diagnosis, and condition indicates whether or not the infant was diagnosed with NEC. kPTR values are provided for cases where there was a clear match with results from the kPTR software^6^ (Supplementary Table 8). Relative abundance was calculated for each organism based on the number of sequencing reads mapped to the genome sequence as a percent of sequences mapped to all draft-quality genomes.

**Supplementary Table 6** Single copy gene inventory for genomes reconstructed as part of this study from a previously published adult human metagenome^16^.

**Supplementary Table 7**iRep measurements for organisms associated with an adult human microbiome^16^.

**Supplementary Table 8** Results from the kPTR software^6^ for the premature infant metagenomes ^20^.

**Supplementary Table 10** iRep measurements for previously sampled Candidate Phyla Radiation (CPR) organisms^12^.

